# Structural basis of Retron-Eco8-mediated anti-phage defense

**DOI:** 10.64898/2025.12.04.692459

**Authors:** Chao Xiong, Huan Pu, Yiwen Tang, Ting Liu, Dantong Luo, Qiang Chen, Yamei Yu

**Affiliations:** Department of Biotherapy, Cancer Center and State Key Laboratory of Biotherapy, West China Hospital, Sichuan University, Chengdu 610041, China

**Author notes:** These authors contributed equally to this work (Chao Xiong, Huan Pu, Yiwen Tang).

## Abstract

Retrons represent a novel class of bacterial defense systems that employ reverse transcriptase (RT), non-coding RNA, and effector proteins to counteract phage infections. In this study, we elucidate the molecular mechanism of a retron system, Retron-Eco8. Biochemical experiments reveal that the Retron-Eco8 holocomplex, rather than the effector alone, exhibits double-stranded DNA cleavage activity in an ATP-independent manner, triggering abortive infection and effectively halting phage propagation. Cryo-electron microscopy (cryo-EM) analysis reveals a supramolecular assembly comprising four RT subunits, four msDNA (multicopy single-stranded DNA) molecules, and four OLD (overcoming lysogenization defect) nucleases—a configuration critical for anti-phage defense. Structural comparisons between apo and ATP-Mg^2+^-bound states demonstrate a local conformational change. Notably, we identify the phage SSB (single-stranded DNA-binding) protein as an activator of Retron-Eco8, and phylogenetic analysis of SSB proteins further elucidates the phage resistance specificity. Collectively, our findings delineate the structural architecture of the Retron-Eco8 defense complex and provide mechanistic insights into retron-mediated bacterial immunity.

## Introduction

The evolutionary arms race between bacteria and bacteriophages has driven the development of diverse and intricate anti-phage defense mechanisms in prokaryotes. While our understanding of these systems is growing, the mechanisms of many remain incompletely characterized^1-3^. The most well-known of these, restriction-modification (RM) and CRISPR-Cas, have been paradigm-shifting, not only for fundamental microbiology but also for their development into indispensable molecular biology tools^4, 5^. Intriguingly, advancing research reveals that core components of prokaryotic immune systems share significant structural and functional homology with elements of eukaryotic innate immunity, suggesting deep evolutionary conservation^6^. Therefore, dissecting bacterial defense mechanisms provides not only invaluable insights into universal immunity principles but also promises a pipeline for developing novel biotechnological tools.

Retrons are a genetic element found in bacteria, typically consisting of a reverse transcriptase (RT) and an adjacent non-coding RNA (ncRNA)^7^. They were first discovered through identification of a multi-copy single-stranded DNA (msDNA) in *Myxococcus xanthus*^*8*^. The ncRNA comprises of two functionally distinct segments, msrRNA and msdRNA (the template for msdDNA), which are flanked by a pair of inverted repeats (IRs, IRa1 and IRa2) located at the 5’ end of the msrRNA and the 3’ end of the msdRNA, respectively^9-14^. The RT recognizes the ncRNA structure and initiates reverse transcription using the 2’-OH group of a conserved guanosine within the msrRNA region as a primer. It then synthesizes msdDNA complementary to the msdRNA segment. During this process, RNase H activity selectively degrades the msdRNA template while preserving non-template regions, resulting in a mature DNA-RNA hybrid^15, 16^.

Leveraging their unique biosynthetic mechanism, retrons have been repurposed as innovative genome-editing tools. Unlike conventional methods, retron-based editing operates independently of DNA double-strand breaks, enabling highly efficient homology-directed repair (HDR) while exhibiting minimal cytotoxicity. These features make retron technology as a powerful and precise platform for genetic manipulation^17, 18^

In recent years, retrons have emerged as anti-phage defense system that function through partnerships with diverse effector proteins, including nucleases, transmembrane signaling proteins and ATPase^19-22^. Structural and mechanistic studies have begun to unravel the operational principles of several retron systems. For examples, Retron-Eco4 and another type-I-A Retron system form supramolecular complexes with the Septu effector, which remain inactive until phage infection triggers complex dissociation to activate the effector to degrade either single-stranded DNA or tRNA, thereby inhibiting phage replication ^12, 14^. In contrast, the Retron-Eco1 system utilizes a nucleoside deoxyribosyltransferase-like effector that polymerizes into filaments to exert anti-phage activity^13, 23^. Another intriguing system is the Retron-Eco8, which is coupled with an effector from the OLD (overcoming lysogenization defect) family, which contains an N-terminal ABC ATPase domain and a C-terminal Toprim nuclease domain. Despite its identification, the mechanism by which Retron-Eco8 mediates anti-phage defense has remained unclear.

In this study, we present structural and biochemical insights into the Retron-Eco8 system. Using Cryo-electron microscopy (cryo-EM), we determined structures of the supramolecular complex in both apo and ATP-Mg^2+^-bound states. These structures reveal a stable assembly with 4:4:4 stoichiometry, comprising msDNA, reverse transcriptase (RT), and the OLD effector—an architecture essential for anti-phage function. Upon phage infection, the complex degrades double-stranded DNA, targeting both phage and *Escherichia coli* genomic DNA, in a manner independent of ATP hydrolysis. Furthermore, we also identify the single-stranded DNA-binding protein (SSB) of phage T3 as an activator of the Retron-Eco8 system. Our findings elucidate a distinct mechanism of retron-mediated immunity and underscore the versatility of retron-effector partnerships in bacterial defense.

## Results

### The Retron-Eco8 system provided defense against phage via abortive infection

The Retron-Eco8 system has been previously demonstrated to play a crucial role in phage defense^19^. To elucidate the molecular mechanism of its anti-phage function, we synthesized the genes of RT, ncRNA, and OLD of the Retron-Eco8 system from *E. coli* 200499 (Fig. 1a), under the control of its native promoter. Then we transferred this system into *E. coli* BL21 (DE3), which naturally lacks this defense system. Subsequently, we challenged the system-containing strain with T phages and observed that the Retron-Eco8 system provided robust protection against T2, T3, T4, T6 and T7, reducing plaque-forming efficiency by 4 to 5 orders of magnitude (Fig. 1b & S1a). Only moderate protection was observed against T5, and no defense was observed against T1 (Fig. 1b & S1a).

**Figure 1.**
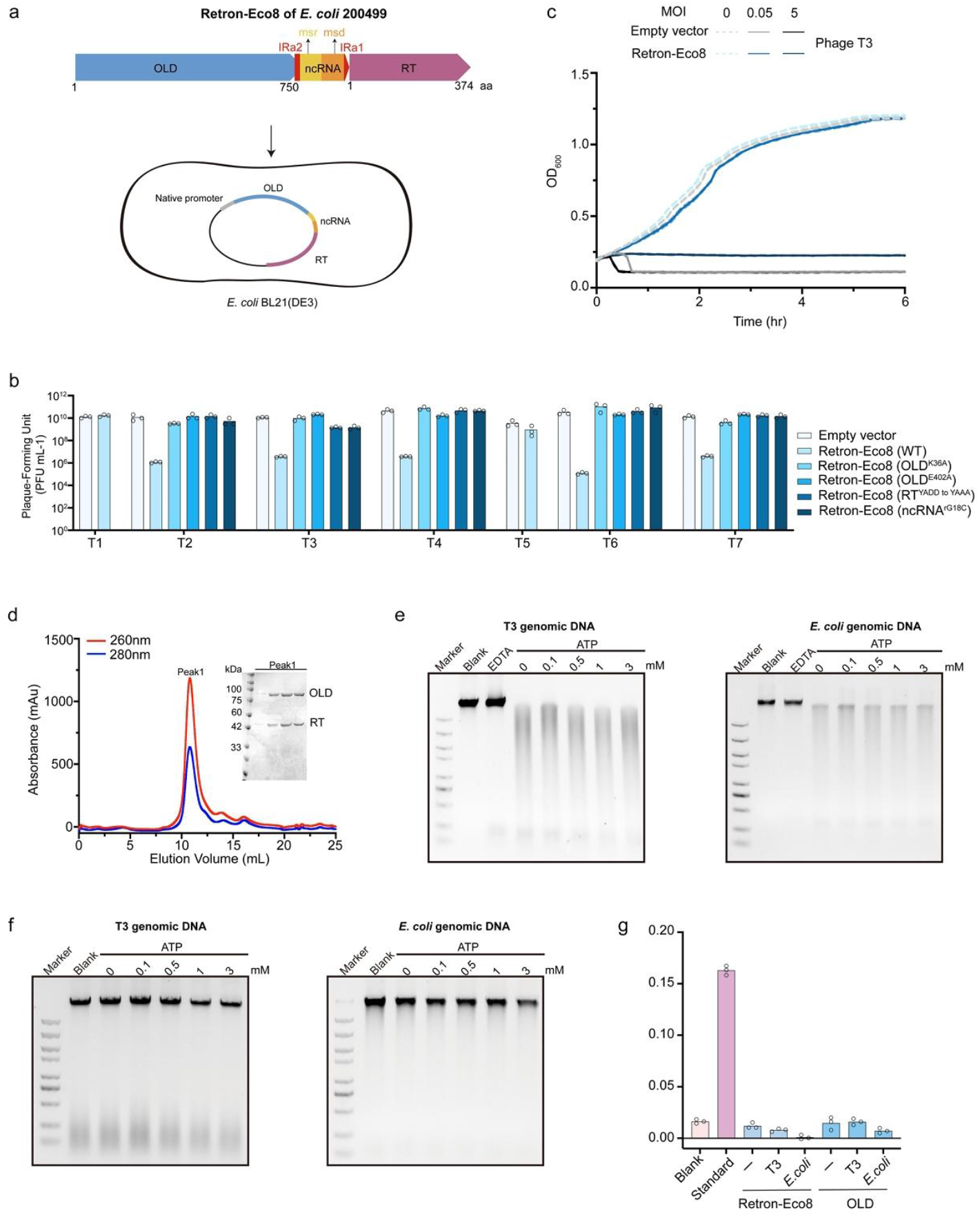
Retron-Eco8 provides defense against phages via abortive infection by nuclease activity. (a) Schematic diagram of Retron-Eco8 system. *E. coli* BL21 (DE3) is used to evaluate the defense activity. (b) Anti-phage activity of Retron-Eco8 system. Plating efficiency of various phages infecting the *E. coli BL21*(DE3) strain containing empty vector, wild-type or mutant Retron-Eco8. Data represent plaque-forming units (PFUs) per milliliter of each phage infection. Bar graphs show the average of three replicates with individual data points overlaid. (c) Phage infection in liquid cultures of the *E. coli* BL21(DE3) strain containing Retron-Eco8. Empty vector is used as the control. Cells are infected at various MOI values. For each MOI, results of three experiments are presented as the average of three replicates with shaded areas indicating SD. (d) Size-exclusion chromatography profiles for the co-expression sample of RT, ncRNA, and OLD. The eluted peak is evaluated by SDS-PAGE. (e) The nuclease activity of Retron-Eco8 holocomplex. (f) The nuclease activity of the individual OLD. (g) The ATPase activity of Retron-Eco8 holocomplex or the individual OLD. The genomic DNA of T3 phage or *E. coli* does not enhance the ATPase activity. 0.5 μM standard phosphorus solution is used as the positive control. Bar graphs show the average of three replicates with individual data points overlaid.

To further dissect the functional requirements, we introduced specific point mutations into critical residues of each component: in OLD, targeting either the ATPase domain (K36A) or the TOPRIM nuclease domain (E402A); in RT, disrupting the catalytic motif YADD^198–201^ by mutating it to YAAA^198–201^; and in ncRNA, mutating the priming guanosine within the msr region to cytosine (rG18C). Each of these mutations abolished the defense of Retron-Eco8 (Fig. 1b). Together, these results demonstrate that the enzymatic activities of RT and OLD, as well as the structural integrity of the ncRNA, are essential for the anti-phage function of this system.

Notably, Retron-Eco8-mediated defense exhibited clear phage load dependence. At low multiplicities of infection (MOI < 1), the system-containing cells maintained vigorous proliferation, whereas high MOI (MOI > 1) triggered rapid cell death—a hallmark feature of abortive infection (Fig. 1c & S1b). Our data demonstrated that Retron-Eco8 provided defense against phages via Abi mechanism, as observed in other retron systems^19^.

### The enzymatic activities of the Retron-Eco8 system

The effector protein of the Retron-Eco8 system belongs to the overcoming lysogenization defect (OLD) family of nuclease. It features an N-terminal ATP-Binding Cassette (ABC) ATPase domain and a C-terminal Topoisomerase/primase (Toprim) domain, with nuclease activity that is metal ion-dependent^24, 25^. We therefore evaluated the nuclease activity associated with this defense system. We co-expressed all elements of the Retron-Eco8 system with a C-terminal His-tag fused to RT. This approach successfully isolated a stable ribonucleoprotein complex (Fig. 1d). *In vitro* assays demonstrated that this complex degraded both phage and bacterial genomic DNA most efficiently in the presence of Mn^2+^ or Co^2+^ (Fig. 1e & S2). To our surprise, *in vitro* assays demonstrated that OLD alone is unable to degrade either phage or bacterial genomic DNA, regardless of ATP presence (Fig. 1f). Over-expression of OLD alone does not demonstrate cytotoxicity (Fig. S3). These findings suggest that OLD’s nuclease activity depends on its assembly into a supramolecular structure of Retron-Eco8.

We further assessed the *in vitro* ATPase activity of Retron-Eco8 complex and the individual OLD. However, neither exhibited detectable ATPase activity under standard assay conditions (Fig. 1g). Given that certain ATPases require DNA substrates for activity^26-30^, we further measured their ATPase activity in the presence of DNA substrates. However, no ATPase activity was observed for either the complex or OLD alone (Fig. 1g).

### RT, msDNA and OLD assemble into a tetrameric supramolecular complex

To understand the structural basis of Retron-Eco8 function, we solved its cryo-EM structure at a resolution of 2.57 Å (Fig. S4). The structure reveals a dodecameric complex with a butterfly-shaped architecture, composed of four copies each of msDNA (containing msrRNA and msdDNA), reverse transcriptase (RT), and the OLD effector, resulting in an overall 4:4:4 stoichiometry (Fig. 2a-c). The msDNA is supposed to be synthesized by RT with the ncRNA as the template. Within this architecture, the OLD effectors form a central tetrameric core, while four RT subunits are positioned at the vertices of the assembly. The msdDNA extends through the RT surface and forms a lasso-like motif encircling two alpha helices from the neighboring OLD molecule, and the msrRNA extensively wraps around the surface of RT (Fig. 2c & 2d).

**Figure 2.**
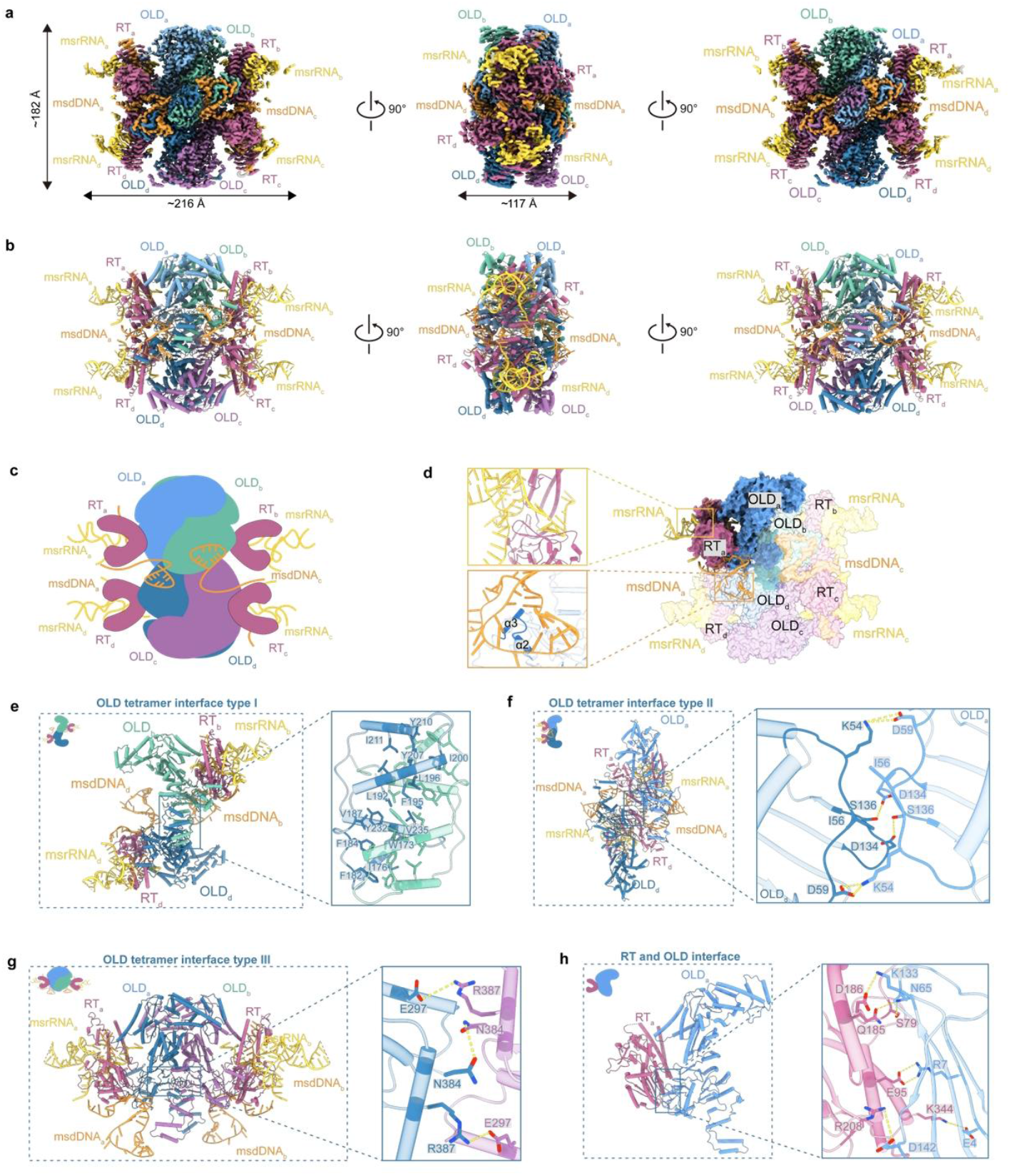
Assembly of Retron-Eco8 supramolecular complex. (a) Cryo-EM density map of Retron-Eco8 complex. (b) Atomic model of Retron-Eco8 complex. (c) Schematic diagram of Retron-Eco8 assembly. (d) Surface representation of Retron-Eco8. One subunit is highlighted by dark colors. The interactions between nucleic acids and proteins are shown in the enlarged part. (e-g) The three types of OLD tetramer interfaces. (h) The interface between RT and OLD.

The central OLD tetramer is assembled through three distinct types of interfaces. Type I interface is formed by the dimerization domains of two diagonal OLD molecules, where four α-helixes from each dimerization domain symmetrically intertwine to create an extensive hydrophobic core (Fig. 2e). Type II and type III interfaces are both mediated by the OLD ATPase domains, establishing the lateral and longitudinal intra-tetramer contacts, respectively (Fig. 2f & 2g).

Each RT molecule binds to an OLD protomer, positioning four RTs peripherally around the OLD tetramer (Fig. 2a-c). The contacts between RT and OLD involve the palm domain of RT and the OLD ATPase domain, primarily stabilized by hydrophilic interactions (Fig. 2h).

### Structure of Retron-Eco8 RT

The RT of Retron-Eco8 adopts a canonical right-hand-like architecture—a hallmark of reverse transcriptases—comprising three subdomains: fingers (residues 41–58), palm (residues 1–40 and 59–266), and thumb (residues 267–374) (Fig. 3a). The overall structure of Retron-Eco8 RT consists of 12 α-helices and 9 β-strands. Specifically, the fingers subdomain contains two β-strands (β1–β2); the palm subdomain encompasses eight α-helices (α1–α8) and two antiparallel β-sheets—one with four strands (β3–β6) and another with three (β7–β9); while the thumb subdomain is composed of four α-helices (α9–α12) (Fig. 3b). Notably, the catalytic YADD motif (residues 198–201) is positioned between strands β4 and β5 in the palm domain (Fig. 3b). The overall topology of RT is highly conserved across diverse retron systems, with well-overlaid YADD motifs (Fig. 3c & 3d). Additionally, the catalytic YADD motif exhibits evolutionary conservation across species (Fig. S5).

**Figure 3.**
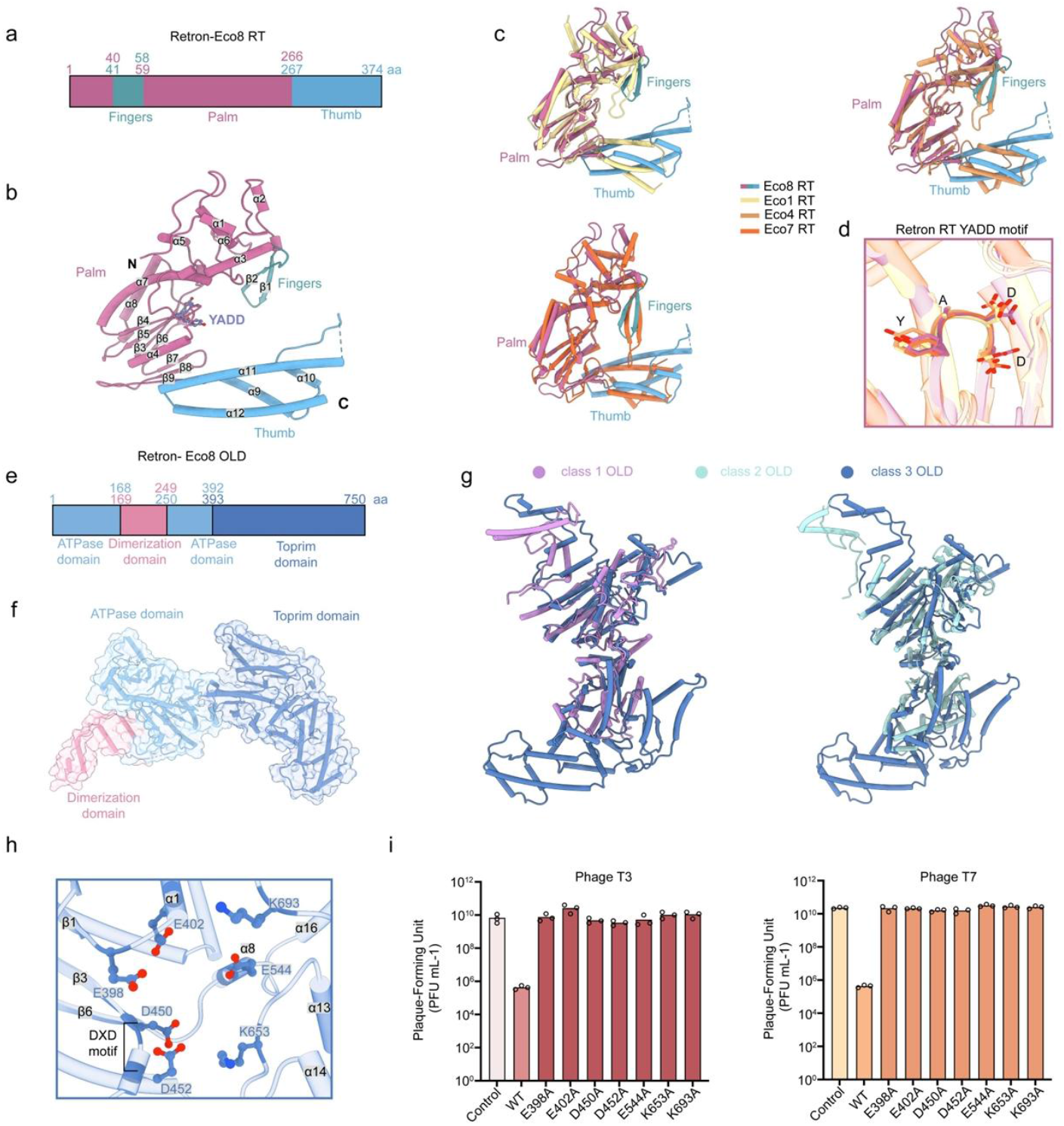
Structures of Retron-Eco8 RT and OLD. (a) Schematic representation showing the domain architectures of Retron-Eco8 RT. (b) Structure of Retron-Eco8 RT. The secondary structures and the catalytic motif YADD are labelled. (c) Superimposition of Retron-Eco8 RT structure with that of Retron-Eco1 (PDB: 7V9U), Retron-Eco4 (PDB: 9E8Z) or Retron-Eco7 (PDB: 9NNB). (d) Superimposition of the catalytic YADD motif of various RTs. (e) Schematic representation showing the domain architectures of Retron-Eco8 OLD. (f) Structure of Retron-Eco8 OLD. (g) Superimposition of the class 3 OLD (Retron-Eco8) structure with that of class 1 (Ts OLD, PDB: 6P74) or class 2 (GajA, PDB: 8SM3). (h) The active site of Retron-Eco8 Toprim nuclease. The conserved catalytic residues are shown as sticks and labelled. (i) Mutations of Toprim active site abolish the Retron-Eco8 defense against T3 and T7.

### Structure of Retron-Eco8 OLD

The effector OLD is comprised three structural domains: an N terminal ATPase domain, a dimerization domain, and a Toprim domain (Fig. 3e & 3f). The ATPase domain consists of 14 α-helices and 11 β-strands, forming a sandwich-like structure in which α1 is flanked by the β-sheets (Fig. S6). Specifically, β-strands 1, 2, 4, 5, and 6 form an antiparallel β-sheet, while strands 3, 7, 8, 9, and 10 constitute a parallel β-sheet. Helices α2–5 and α10–14 encircle this β-sheet core (Fig. S6). The dimerization domain, composed of four α-helices (α6–9), inserts between the β7 strand and α10 helix of the ATPase domain and is essential for tetramer assembly (Fig. S6). The Toprim domain contains a four-stranded parallel β-sheet (β1–β2–β3–β6) surrounded by α-helices, with three additional α-helices and two short β-strands inserted between β3 and β6 (Fig. S6). Retron-Eco8 OLD represents the first structure of the class 3 OLD family, exhibiting a significantly expanded Toprim domain augmented with additional α-helices when compared to class 1 and class 2 OLD structures (Fig. 3g). This architectural divergence implies its distinct evolutionary trajectory within the OLD protein superfamily.

The nuclease activity of the effector OLD is performed by the Toprim domain, which is supposed to adopt a two-metal mechanism for nuclease cleavage^24, 25^. The Toprim active site comprises a canonical glutamate residue E398 located immediately after β1, a DXD motif situated between β3 and β4, two additional conserved glutamate residues E402 and E544 in α1 and α8, and two positively charged residues K653 (between α13 and α14) and K693 (between α16 and α17) (Fig. 3h). Evolutionary analysis confirms that these residues are highly conserved across species harboring the Retron-Eco8 system (Fig. S7). Point mutations in any of these catalytic residues completely abolish anti-phage activity against T3 and T7 phages (Fig. 3i), underscoring the essential role of nuclease activity in phage defense.

### Structure of msDNA

Like other characterized retron systems^12-14^, the ncRNA of the Retron-Eco8 is organized into four regions: IRa2, IRa1, msrRNA and msdRNA (Fig.4a). The Retron-Eco8 RT initiates at a highly conserved guanine residue on the ncRNA and uses it as template to synthesis of msDNA *in vivo* (Fig. S8). Although the secondary structure of Retron-Eco8 msDNA has not been experimentally determined yet, the high-resolution cryo-EM density maps enabled us to unambiguously identify the nucleobases of the msDNA and resolve its three-dimensional structure (Fig. 4b & 4c). In the resolved structure, IRa1, IRa2 and DSLb were not visible, while the msrRNA (rA19-rG81) and most of the msdDNA (dC1-dT35, dC66-dT75) are clearly observed (Fig. 4c). A unique 2′,5′-phosphodiester bond links the branching guanine (rG18 in msrRNA) and the initiating dedeoxycytidine (dC1 in msdDNA) (Fig.4b & 4c). For the Retron-Eco8 msDNA, the msrRNA contains two stem loops (RSLa, rA28-rU45, and RSLb, rA46-rU61), four single-stranded RNA segments, and one DNA-RNA duplex between msrRNA (rA73-rC78) and msdDNA (dG70-dT75). The msdDNA consists of three single-stranded DNA regions (ssDa, dC1-dT10, ssDb, dT35-dA37, ssDc, dA62-dA69), two DNA stem loops (DSLa, dC11-dG34, and DSLb, dC38-dG61) and one DNA-RNA duplex (Fig.4b & 4c). The two DNA stem loops is unique to Retron-Eco8, since the reported Retron-Eco1, Retron-Eco4, and Retron-Eco7 contain only one DNA stem loop^12-14^.

**Figure 4.**
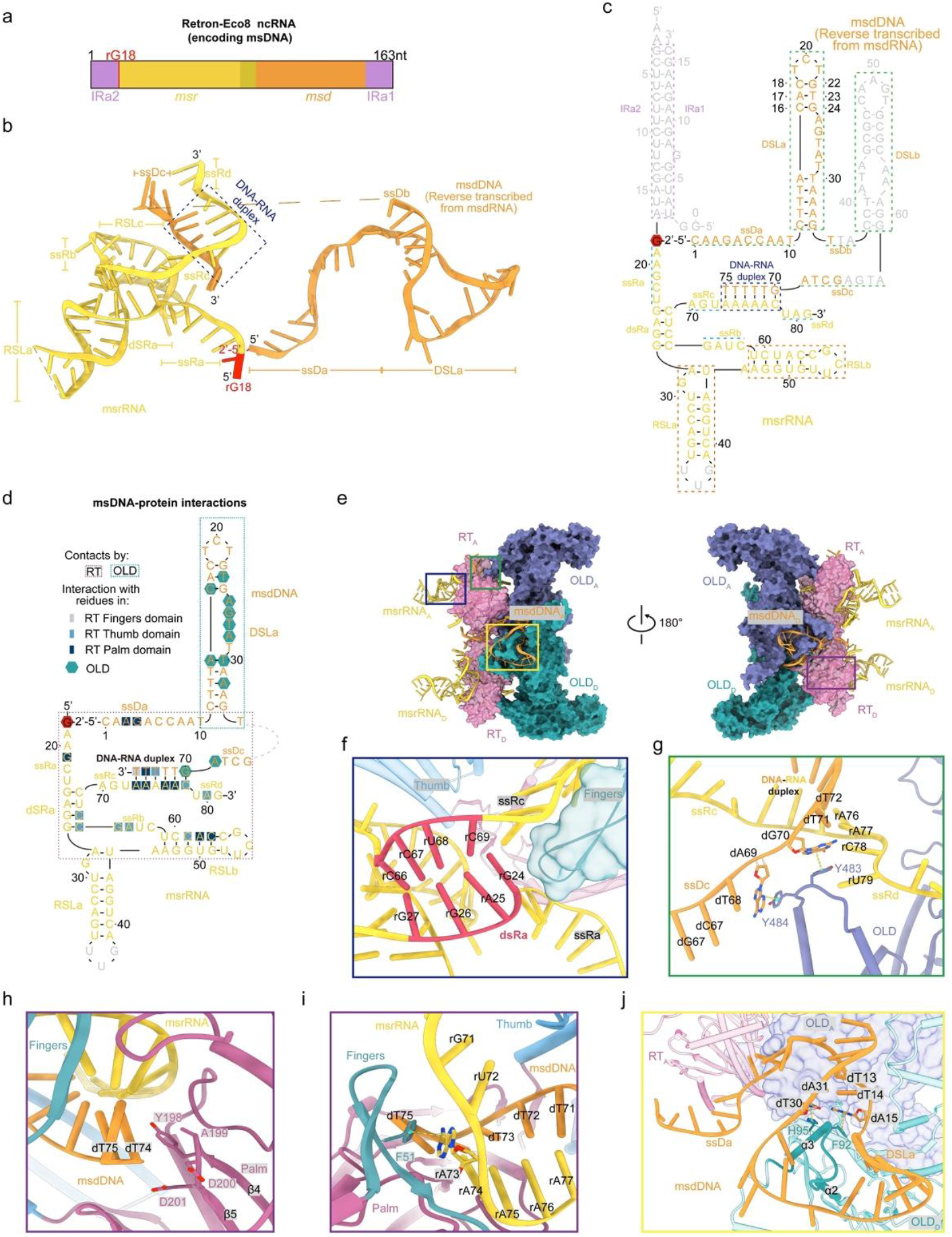
Interactions between msDNA and the proteins. (a) Schematic representation of Retorn-Eco8 ncRNA. (b) Solved tertiary structure of Retron-Eco8 msDNA. (c) Secondary structure of Retron-Eco8 msDNA. The DNA–RNA duplex and stem loops are indicated by dotted boxes. The regions not observed in the solved structure are colored grey. (d) Schematic representation of msDNA showing an overview of its interaction with RT and OLD. (e) The msDNA interacts with both RT and OLD. For clarity, only two subunits are shown. (f) RT fingers subdomain sterically disrupts the double-stranded RNA. (g) OLD Y483 and Y484 pack against dG70 and dA69, mediating the branching of ssDc and ssRd. (h) The catalytic YADD motif is located close to the 3’ end of the msdDNA. (i) RT F51 paired with rA73. (j) The msdDNA interacts with both RT and the adjacent OLD. F92 and H95 pack against dA15 and dT30, respectively, facilitating the formation of the lasso-like configuration.

In the cryo-EM structure of Retron-Eco8, the DNA stem loop DSLa exhibits 8 base pairs, in contrast to the 10 base pairs suggested by its predicted secondary structure (Fig. S9). Despite the predicted structure being energetically more favorable, the observed structure of DSLa allows the formation of a 5-nucleotide lasso-like loop. This loop encircles the two α-helices of an adjacent OLD molecule, thereby promoting the assembly of the compact Retron-Eco8 complex (Fig. 2d).

### msDNA engages in interactions with both RT and OLD to stabilize the supramolecular complex

Structural analysis reveals that the msDNA wraps around the electropositive surfaces of both RT and OLD (Fig. S10). Extensive intermolecular contacts are observed between the msDNA and both RT and OLD, while the msrRNA predominantly engages with RT and the msdRNA primarily interacts with OLD (Fig. 4d).

The interactions between msrRNA and RT are mediated by the ssRa, ssRb, dsRa and RSLb segments, while RSLa extends outward and does not participate in direct contacts with the proteins (Fig. 4d & 4e). At one end of dsRa, the fingers subdomain of RT sterically disrupts the double-stranded RNA, driving its separation into two single-stranded segments (ssRa and ssRc) (Fig. 4f). The DNA-RNA duplex (rA73-rC78 paired with dG70-dT75) is positioned within the central cavity of the RT, embraced by the thumb and fingers subdomains. Two aromatic residues Y483 and Y484 of OLD pack against dG70 and dA69, mediating the branching of ssDc and ssRd (Fig. 4g). The 3′ end of the msdDNA (T75) is positioned directly adjacent to the YADD active motif (Fig. 4h). F51 of the fingers subdomain packs against the base of rA73, which pairs with dT75 (Fig. 4i). This interaction likely sterically blocks further extension of msdDNA, thereby enabling RT termination in retron-Eco8.

The ssDa segment of the msdDNA wraps around the RT surface, and the DSLa segment of the msdDNA forms a lasso-like configuration encircling two alpha helices (α2 and α3) from the neighboring OLD molecule (Fig. 4j). Two ring-residues F92 and H95 of the encircled α3 pack with dA15 and dT30, respective (Fig. 4j). Thus, msdDNA ties up the two adjacent subunits, playing an essential role in facilitating the assembly of the Retron-Eco8 supramolecular complex.

### Structure of Retron-Eco8 complexed with ATP

To provide the structural insights into the functional role of ATP for the Retron-Eco8 function, we determined the cryo-EM structure of the Retron-Eco8 complexed with ATP– Mg^2+^ at 2.8 Å resolution (Fig. 5a & S4). Well-defined electron density for ATP reveals a complete nucleotide-binding mode within the OLD ATPase domain of each protomer (Fig. 5a). The ATPase domain adopts a canonical ABC ATPase fold, featuring six conserved motifs: Walker A, Walker B, signature motif, Q-loop, D-loop, and H-loop (Fig. S11). Notably, the four ATP molecules are located at the central cavity of the OLD tetramer (Fig. 5a & 5b). ATP binding induced ordering of a previously disordered loop (V242–G255) of the adjacent OLD’s ATPase domain (Fig. 5b). Despite these local rearrangements, structural alignment between the apo and ATP-bound OLD tetramers revealed only minimal global conformational changes, with a root mean square deviation (RMSD) of 0.69 Å for 747 Cα atoms (Fig. 5b), indicating that ATP binding does not substantially alter the tetrameric architecture.

**Figure 5.**
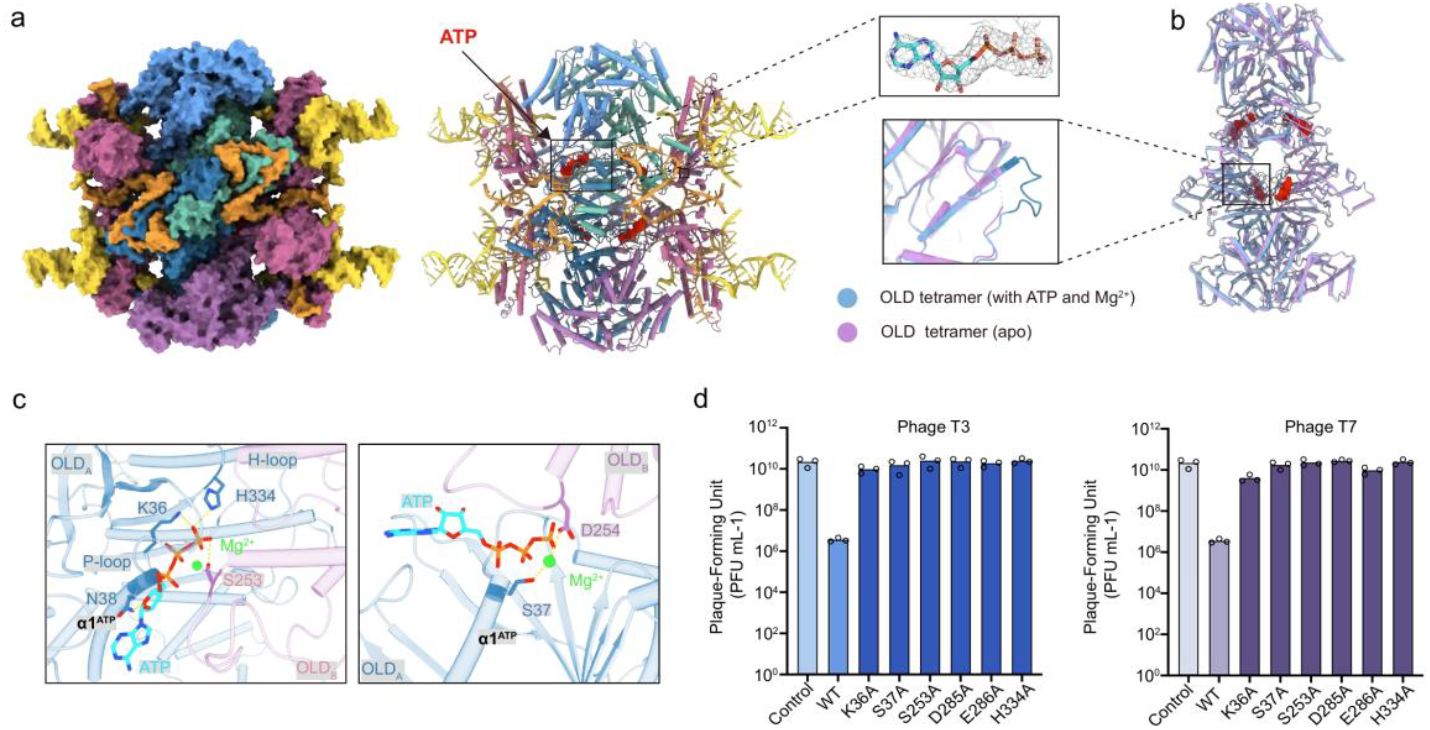
Structure of Retron-Eco8 complexed with ATP–Mg^2+^. (a) Overall structure of ATP-bound Retron-Eco8. Left, cryo-EM map; Right: atomic model. The density map of ATP is colored grey and contoured at 3.0 σ. (b) Overlayed structures of OLD tetramer in apo and ATP-bound states. The ATP molecules are shown in sphere. The ATP-induced ordered loop is shown in the enlarged part. (c) The ATP binding mode in OLD. (d) Mutations of ATPase active site abolish the Retron-Eco8 defense against T3 and T7.

In the ATP-binding pocket, the triphosphate moiety of ATP is coordinated by the Walker A motif (residues 30–38) and H334, while Mg^2+^ is chelated by the β- and γ-phosphates of ATP, S37 of P-loop and D254 of the adjacent OLD’s ATPase domain (Fig. 5c). Point mutations in any of these key residues completely abolish the Retron-Eco8 anti-phage activity against T3 and T7 phages (Fig. 5d), indicating that the ATP-binding plays some essential role in phage defense.

### SSB is a phage trigger for Retron-Eco8

A previous study has suggested that phage single-stranded DNA-binding protein (SSB) acts as a trigger for the Retron-Eco8 defense system^20^. Our plaque assays demonstrate that the Retron-Eco8 system confers resistance against multiple T-phages except T1 (Fig. 1b). If SSB could activate Retron-Eco8, co-expression of SSB and Retron-Eco8 will result in cell death. We then sought to determine whether SSB proteins from various T-phages could activate Retron-Eco8. Over-expression of SSB alone from T2, T4, or T6 phage induces strong cellular toxicity, which precluded definitive assessment of their ability to activate Retron-Eco8 (Fig. S12). Co-expression of Retron-Eco8 with SSB from T3 or T7 results in severe cytotoxicity, while moderate cytotoxicity is observed for T5 SSB and no cytotoxicity is observed for T1 or *E. coli* SSB (Fig. 6a). Bacterial growth curve analyses confirm that T3 or T7 SSB triggers Retron-Eco8-mediated cell death (Fig. 6b).

**Figure 6.**
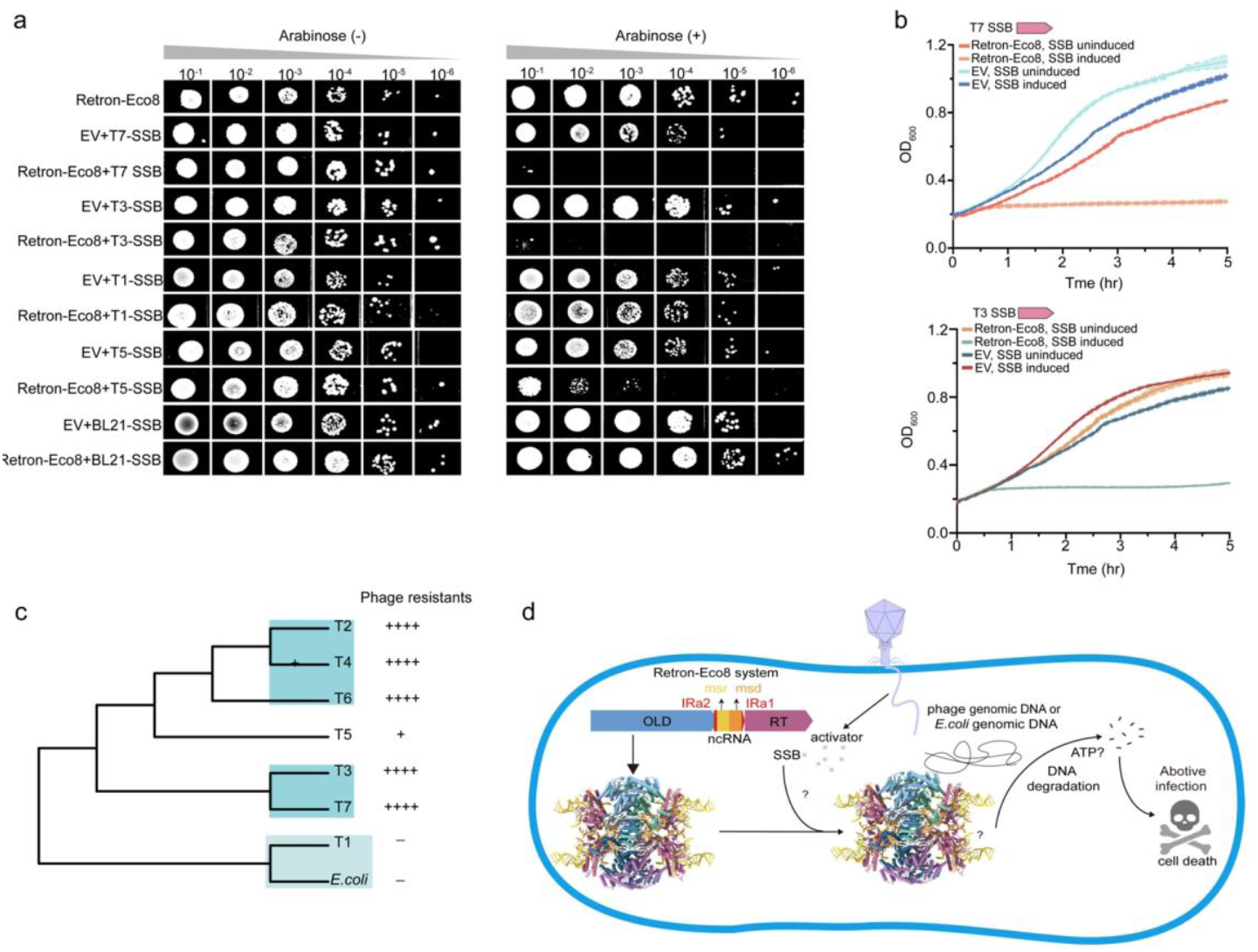
Phage SSB is a trigger for the Retron-Eco8 system. (a) Representative plating assay shows the cytotoxicity of co-expression of Retron-Eco8 and SSB. (b) Growth curves for *E. coli* BL21(DE3) cells show that co-expression of Retron-Eco8 and SSB results in cell death. Results of three experiments are presented as the average of three replicates with shaded areas indicating SD. (c) Maximum-likelihood phylogenetic tree of SSB proteins from T-phages and *E. coli*. (d) A proposed model of anti-phage defense of the Retron-Eco8 system.

SSB is a highly conserved protein present in diverse organisms, including the host *Escherichia coli*. To avoid autoimmunity, *E. coli* SSB must not activate Retron-Eco8, as demonstrated by the cell-toxicity assays (Fig. 6a). Our phylogenetic analysis of SSB proteins from T-phages and their host, *E. coli*, reveals a clear evolutionary basis for phage resistance. The SSB of the resistant T1 phage clusters within the same clade as the host SSB, distinctly separate from the clade containing SSBs of Retron-Eco8-sensitive phages (Fig. 1b & 6c). This distinct clustering suggests that T1 SSB’s sequence similarity to *E. coli* SSB prevents its recognition by the Retron-Eco8 system (Fig. 1b & 6c).

## Discussion

Retrons, which consist of a reverse transcriptase (RT) and an adjacent non-coding RNA (ncRNA), form antiviral defense systems by coupling with diverse effector proteins to implement Abi strategies against phages^2, 7, 19^. In this study, we employed cryo-EM to determine high-resolution structures of Retron-Eco8 in both apo and ATP-Mg^2+^-bound states, revealing a 4:4:4 stoichiometry of msDNA, RT, and the OLD effector (Fig. 2 & 5). Biochemical assays established that the nuclease activity of Retron-Eco8 is dependent on its assembly into a supramolecular complex (Fig. 1e & 1f).

Retron-Eco4 and Retron-Eco7 employ PtuAB as the effector, and phage invasion induces msDNA degradation via either PtuAB or phage nuclease and prompts dissociation of the complex to release the effector nuclease resulting in subsequent DNA degradation and phage resistence^12, 14^. Retron-Eco8 employs a class 3 OLD as its effector. In contrast to the reported retron systems, Retron-Eco8 exhibited its nuclease activity via holocomplex, rather than the effector alone (Fig. 1e & 1f). The effector OLD alone performed neither cytotoxicity *in vivo* nor nuclease activity *in vitro* (Fig. S3 & 1f).

The OLD family of nucleases is characterized by an N-terminal ATPase domain and a C-terminal topoisomerase/primase (Toprim) domain^31, 32^. Based on local gene neighborhood relationships, OLD homologs could be clustered into three groups^33^. Class 1 OLD comprises standalone genes, exemplified by Ts OLD and P2 OLD^24, 34^. Class 2 OLD is synonymous with GajA, which operates in conjunction with GajB to form the multi-component defense system Gabija^1^. Class 3 OLD encompasses effector proteins integrated into retron systems, such as Retron-Eco8^19^. Notably, the ATPase activity of OLD enzymes exhibits significant variability, with ATP exerting divergent regulatory effects on their DNA cleavage activity—either enhancing, inhibiting, or leaving it unaffected^35^. The OLD effector in Retron-Eco8 exhibited no detectable ATPase activity either alone or when complexed with RT and msDNA (Fig. 1g). The *in vitro* nuclease activity of Retron-Eco8 seemed ATP-independent (Fig. 1e). However, mutations of the key residues involved in ATP binding or hydrolysis completely abolished the anti-phage activity of Retron-Eco8 (Fig. 5d). Further studies are needed to elucidate how the ATPase domain of the OLD effector contributes to the anti-phage defense of the Retron-Eco8 system.

We identified phage SSB as the trigger for Retron-Eco8 activation, as SSB induced Retron-Eco8-mediated cell death (Fig. 6). To prevent autoimmunity, Retron-Eco8 needs to discriminate self (host SSB) and non-self (phage SSB). Phylogenetic analysis of SSB proteins reveals that SSBs of Retron-Eco8-resistant T1 phage and the host *E. coli* SSB are clustered into a distinct clade separated from those of Retron-Eco8-sensitive phages (Fig. 6c). This genetic distance provides an explanation for the phage-resistance of Retron-Eco8 and guarantees that *E. coli* SSB could not activate Retron-Eco8 defense system thus to prevent autoimmunity.

## Methods

### Bacterial strains and culture conditions

*Escherichia coli* BL21(DE3) was used as the host for all experiments. Cells were routinely cultured in Lysogeny Broth (LB) at 37°C with shaking at 220 rpm. For plasmid maintenance, the medium was supplemented with the following antibiotics as appropriate: ampicillin (100 µg mL^-1^), kanamycin (50 µg mL^-1^), streptomycin (50 µg mL^-1^), or chloramphenicol (34 µg mL^-1^).

### Plasmid construction

The native Retron-Eco8 operon from *E. coli* 200499 was synthesized and cloned into a pACYCDuet-1 vector for functional studys. For heterologous expression, the genes encoding the OLD, ncRNA, and RT were cloned into a pET28a vector, generating a C-terminal 6×His tag on the RT (pET28a-OLD-ncRNA-RT-His_6_). Additionally, the ncRNA was cloned into a pCDFDute-1 vector (pCDFDute-1-ncRNA), and the OLD protein alone was inserted into a pET28a vector with a C-terminal 6×His tag (pET28a-OLD-His_6_). Separately, SSB genes from various phages and *E. coli* were cloned into a pBAD vector. All mutants were generated using Gibson Assembly, with the wild-type construct serving as the template.

### Protein expression and purification

For complex expression, the plasmids pET28a-OLD-ncRNA-RT-His_6_ and pCDFDute-1-ncRNA were co-transformed into *E. coli* BL21 (DE3) cells. Protein expression was induced with 200 µM IPTG at an OD_600_ of ∼0.8. Cells were then incubated at 18°C for 16 hours.

Cells were harvested by centrifugation and lysed in buffer (25 mM Tris-HCl pH 8.0, 500 mM NaCl, 20% glycerol) using a high-pressure homogenizer. The lysate was clarified by centrifugation, and the supernatant was applied to a Ni^2+^-NTA column. The column was washed sequentially with lysis buffer containing 10 mM and 20 mM imidazole. The target protein was then eluted with buffer containing 250 mM imidazole. A final polishing step was performed using size-exclusion chromatography on a Superdex 200 10/300 GL column (GE Healthcare) pre-equilibrated with 25 mM Tris-HCl pH 8.0, 250 mM NaCl, 1 mM DTT. Peak fractions containing the target protein were pooled for subsequent structural and biochemical studies. The OLD protein alone was expressed and purified using the same protocol.

### Phage cultivation and plaque assays

Phages were acquired from DSMZ (T2, T4, T5 and T7) and BNCC (T1, T3 and T7). To propagate, a single plaque was picked and inoculated into a liquid culture of *E. coli* BL21(DE3) at an OD_600_ of 0.2–0.4 and incubated at 37°C until complete lysis occurred. Lysates were clarified by centrifugation and filtration (0.22 µm).

For defense system evaluation, overnight cultures of BL21(DE3) harboring the pACYCDuet-1 vector (encoding the wild-type Retron-Eco8 system, mutants, or empty vector control) were diluted 1:100 in fresh LB with antibiotics and grown to an OD_600_ of 0.2–0.3. 2-mL aliquot was mixed with 20 mL of LB containing 0.6% soft agar and poured over LB agar plates. After the overlay solidified, 3 µL of serial (10-fold) phage dilutions were spotted onto the plates. Plates were incubated at 37°C for 4–6 hours (T1, T3, T7) or overnight (T2, T4, T5, T6). Plaque-forming units (PFU) were counted, and the efficiency of plating (EOP) was calculated as the ratio of PFU on the test strain to PFU on the control strain.

### Phage-infection dynamics in liquid culture

*E. coli* BL21(DE3) cells harboring either the pACYCDuet-1-Retron-Eco8 or an empty pACYCDuet-1 vector (negative control) were grown to early-log phase (OD_600_ ∼0.3). For infection, bacterial cultures were seeded into 96-well plates at 180 µL per well. Subsequently, 20 µL of phage lysate was added to each well to establish final multiplicity of infection (MOI) values of 5 and 0.05, representing high and low infection conditions, respectively. Wells containing LB medium instead of phage lysate were included as uninfected controls. OD_600_ was monitored every 5 minutes for 12 hours using a TECAN Sunrise™ Microplate Reader.

### Toxicity assays on solid/liquid media

To assess whether phage SSB could activate the Retron-Eco8 system and induce cytotoxicity, we performed a co-transformation assay in *E. coli* BL21(DE3). The pBAD plasmid, carrying an SSB gene under the control of an arabinose-inducible araBAD promoter, was co-transformed with the pACYCDuet-1 plasmid harboring the Retron-Eco8 system under its native promoter. Cells carrying the SSB gene or the Retron-Eco8 system alone served as negative controls.

For the CFU-based spot assay, overnight cultures were serially diluted, and 3 µL of each dilution was spotted onto LB agar plates with or without 0.2% arabinose inducer. Following overnight incubation at 37°C, the resulting colonies were imaged and quantified. For real-time growth curve analysis, early-log phase cultures were aliquoted into a 96-well plate. SSB expression was induced by adding arabinose to a final concentration of 0.2%. The growth kinetics were monitored by measuring the OD_600_ every 5 minutes at 37°C with shaking using a TECAN Sunrise™ Microplate Reader.

### Nuclease activity assay

Nuclease activity assays were performed using either the purified OLD protein alone or the reconstituted Retron-Eco8 complex (RT, OLD and msDNA). Standard 10 µL reactions contained approximately 200 ng of genomic DNA, 500 nM of the specified protein, and reaction buffer (25 mM Tris-HCl, 200 mM NaCl, and 10 mM divalent metal chloride). Reactions were incubated for 60 minutes at temperatures ranging from 16°C to 37°C, in the presence or absence of ATP at various concentrations. Reactions were terminated by adding EDTA to 10 mM, followed by Proteinase K treatment to digest proteins. The DNA degradation products were then resolved by 1% agarose gel electrophoresis and visualized.

### ATPase activity assay

We quantified the ATP hydrolysis activity of the Retron-Eco8 complex using a commercial ATPase Assay Kit (Solarbio). Reactions containing 500 nM Retron-Eco8 complex, with or without approximately 200 ng of genomic DNA from various phages or E. coli. The release of inorganic phosphate (Pi) was monitored by measuring absorbance at 660 nm using a SpectraMax® Absorbance Plus Microplate Reader (Molecular Devices).

### Cryo-EM sample preparation and data collection

#### Apo Retron-Eco8

For cryo-EM grid preparation, 3 µL of the protein sample (4.5 mg/mL) was applied to a glow-discharged Quantifoil R2/1 200-mesh Cu grid (Electron Microscopy Sciences). The grid was blotted for 3 seconds at 100% humidity at 4°C and then rapidly frozen in liquid ethane using a Vitrobot Mark IV (Thermo Fisher). A total of 7,920 micrographs were collected using a 300 kV Titan Krios electron microscope (FEI) equipped with K2 Summit camera (Gatan) with a magnification of 165,000×, and a physical pixel size of 0.85 Å with defocus values ranging from −1.0 to −1.8 μm. A total electron dose of 56.58 e^-^/Å^2^ over 30 frames during a total exposure time of 6 s.

#### Retron-Eco8 complexed with ATP-Mg^2+^

To form the complex, the Retron-Eco8 sample (4.5 mg/mL) was incubated with 3 mM ATP and 5 mM MgCl_2_ on ice for 30 minutes prior to grid preparation. Grids were prepared and frozen under identical conditions as the apo sample. A total of 4,331 images were collected using EPU automated software on a Titan Krios G4 transmission electron microscope (Thermo Fisher Scientific), equipped with a Falcon 4i detector and a Selectris energy filter (slit width 10 eV). Data were recorded at a nominal magnification of 165,000× (physical pixel size 0.75 Å) with a total dose of 50 e^–^/Å^2^ in EER format (50 fractions).

### Cryo-EM data processing

All data processing was performed in cryoSPARC v4.1. Motion correction and CTF estimation were performed on the collected movies. Initial particles were picked with blob picking. Maps were post-processed using DeepEMhancer.

For apo Retron-Eco8, the collected micrograph frames were subjected to motion correction using MotionCor2 in RELION 3.1, followed by contrast transfer function (CTF) estimation with patch CTF estimation in cryoSPARC v4.1. A total of 4,013,724 particles were extracted with a box size of 480 pixels for 2D classification. After two rounds of 2D classification, 396,585 particles were selected for multi ab-initio reconstruction and heterogeneous refinement. The best class, containing 303,229 particles, was further refined though homogeneous refinement and non-uniform refinement and local refinement with C2 symmetry, resulting in a final density map with a resolution of 2.57 Å.

For Retron-Eco8 complexed with ATP-Mg^2+^, a total of 2,171,015 particles were extracted with a box size of 360 pixels for 2D classification. After two rounds of 2D classification, 225,512 particles were selected for ab-initio reconstruction and heterogeneous refinement. The best class was further refined through homogeneous refinement, non-uniform refinement, and local refinement, yielding a final density map at 2.8 Å resolution.

### Model building and refinement

The alphaFold2^36^ models for RT and OLD were fitted into the cryo-EM maps using Chimera^37^. Owing to the high resolution, the msDNA could be built de novo into the electron density. The model was then manually rebuilt in Coot^38^ and refined in PHENIX^39^ with real-space refinement, incorporating secondary structure and geometry restraints. The model was validated by MolProbity^40^. All structural figures were prepared using ChimeraX-1.5^41^ and PyMOL (https://pymol.org)

## Acknowledgements

Financial support for this work was provided by National Key R&D Program of China (2025YFC3408400), National Natural Science Foundation of China (32270761), Sichuan Foundation for Distinguished Young Scholars (25NSFJQ0249), Sichuan Science and Technology Program (2024NSFTD0029) and Sichuan Science and Technology Program (25QNJJ4988).

## Author contributions

Q.C. and Y.Y. conceived and designed the experiments. C.X., H.P., Y.T., T.L, D.L. and Y.Y. performed experiments. C.X., Q.C. and Y.Y. analyzed the data and wrote the manuscript.

## Data and code availability

The cryo-EM density maps of Retron-Eco8 in its apo and ATP-Mg^2+^-bound states have been deposited in the Electron Microscopy Data Bank (EMDB) under accession code EMDB-66659 and EMDB-66663, respectively. The atomic coordinates of Retron-Eco8 in its apo and ATP-Mg^2+^-bound states have been deposited in the Protein Data Bank (PDB) under the accession code 9X94 and 9X9B, respectively.

## References

1. Doron S, et al. Systematic discovery of antiphage defense systems in the microbial. pangenome. Science 359, (2018).

2. Gao L, et al. Diverse enzymatic activities mediate antiviral immunity in prokaryotes. Science. 369, 1077–1084 (2020).

3. Millman A, et al. An expanded arsenal of immune systems that protect bacteria from phages. Cell Host Microbe 30, 1556–1569 e1555 (2022).

4. Roberts RJ, Vincze T, Posfai J, Macelis D. REBASE--a database for DNA restriction and modification: enzymes, genes and genomes. Nucleic Acids Res 43, D298–299 (2015).

5. Pacesa M, Pelea O, Jinek M. Past, present, and future of CRISPR genome editing technologies. Cell 187, 1076–1100 (2024).

6. Wein T, Sorek R. Bacterial origins of human cell-autonomous innate immune mechanisms. Nat Rev Immunol 22, 629–638 (2022).

7. Temin HM. Retrons in bacteria. Nature 339, 254–255 (1989).

8. Inouye Tytfsim. Multicopy single-stranded DNA isolated from a gram-negative bacterium, Myxococcus xanthus. Cell 84, 90541–90545 (1984).

9. Inouye Adbltfmis. Structure of msDNA from Myxococcus xanthus: evidence for a long, self-annealing RNA precursor for the covalently linked, branched RNA. Cell 51, 1105–1112 (1987).

10. Inouye MIS. msDNA and bacterial reverse transcriptase. Annu Rev Microbiol 45, 163–168 (1991).

11. Lampson BC, Inouye, M. & Inouye, S. Retrons, msDNA, and the bacterial genome. Cytogenet Genome Res 110, 491–499 (2005).

12. George JT, et al. Structural basis of antiphage defence by an ATPase-associated reverse. transcriptase. Nat Commun 16, 8459 (2025).

13. Wang Y, et al. Cryo-EM structures of Escherichia coli Ec86 retron complexes reveal architecture and defence mechanism. Nat Microbiol 7, 1480–1489 (2022).

14. Wang C, et al. Disassembly activates Retron-Septu for antiphage defense. Science 389, eadv3344 (2025).

15. Inouye Bclmis. Reverse transcriptase with concomitant ribonuclease H activity in the cell-free synthesis of branched RNA-linked msDNA of Myxococcus xanthus. 56, 701–707 (1989).

16. BC Lampson SIM Inouye. msDNA of bacteria. Prog Nucleic Acid Res Mol Biol 40, 1–24 (1991).

17. Kong X, et al. Precise genome editing without exogenous donor DNA via retron editing system in human cells. Protein Cell 12, 899–902 (2021).

18. Liu W, et al. Retron-mediated multiplex genome editing and continuous evolution in. Escherichia coli. Nucleic Acids Res 51, 8293–8307 (2023).

19. Millman A, et al. Bacterial Retrons Function In Anti-Phage Defense. Cell 183, 1551-1561. e1512 (2020).

20. Stokar-Avihail A, et al. Discovery of phage determinants that confer sensitivity to bacterial. immune systems. Cell 186, 1863–1876 e1816 (2023).

21. Bobonis J, et al. Bacterial retrons encode phage-defending tripartite toxin-antitoxin systems. Nature 609, 144–150 (2022).

22. Mestre MR, Gonzalez-Delgado A, Gutierrez-Rus LI, Martinez-Abarca F, Toro N. Systematic prediction of genes functionally associated with bacterial retrons and classification of the encoded tripartite systems. Nucleic Acids Res 48, 12632–12647 (2020).

23. Carabias A, et al. Retron-Eco1 assembles NAD(+)-hydrolyzing filaments that provide. immunity against bacteriophages. Mol Cell 84, 2185–2202 e2112 (2024).

24. Schiltz CJ, Adams MC, Chappie JS. The full-length structure of Thermus scotoductus OLD defines the ATP hydrolysis properties and catalytic mechanism of Class 1 OLD family nucleases. Nucleic Acids Res 48, 2762–2776 (2020).

25. Schiltz CJ, Lee A, Partlow EA, Hosford CJ, Chappie JS. Structural characterization of Class 2 OLD family nucleases supports a two-metal catalysis mechanism for cleavage. Nucleic Acids Res 47, 9448–9463 (2019).

26. Rzechorzek NJ, Blackwood JK, Bray SM, Maman JD, Pellegrini L, Robinson NP. Structure of. the hexameric HerA ATPase reveals a mechanism of translocation-coupled DNA-end processing in archaea. Nat Commun 5, 5506 (2014).

27. Yoder BL, Burgers PM. Saccharomyces cerevisiae replication factor C. I. Purification and characterization of its ATPase activity. Journal of Biological Chemistry 266, 22689–22697 (1991).

28. Massey TH, Mercogliano CP, Yates J, Sherratt DJ, Lowe J. Double-stranded DNA translocation: structure and mechanism of hexameric FtsK. Mol Cell 23, 457–469 (2006).

29. Yang J, Sun Y, Wang Y, Hao W, Cheng K. Structural and DNA end resection study of the. bacterial NurA-HerA complex. BMC Biol 21, 42 (2023).

30. Shen BW, et al. Structure, substrate binding and activity of a unique AAA+ protein: the BrxL phage restriction factor. Nucleic Acids Res 51, 3513–3528 (2023).

31. Koonin EV GA. The superfamily of UvrA-related ATPases includes three more subunits of putative ATP-dependent nucleases. Protein Seq Data Anal 5, 43–45 (1992).

32. Aravind L LD, Koonin EV. Toprim--a conserved catalytic domain in type IA and II topoisomerases, DnaG-type primases, OLD family nucleases and RecR proteins. Nucleic Acids Res 26, 4205–4213 (1998).

33. Dot EW, Thomason LC, Chappie JS. Everything OLD is new again: How structural, functional, and bioinformatic advances have redefined a neglected nuclease family. Mol Microbiol 120, 122–140 (2023).

34. Myung H CR. The old exonuclease of bacteriophage P2. J Bacteriol 177, 497–501 (1995).

35. Akritidou K, Thurtle-Schmidt BH. OLD family nuclease function across diverse anti-phage defense systems. Front Microbiol 14, 1268820 (2023).

36. Jumper J, et al. Highly accurate protein structure prediction with AlphaFold. Nature 596, 583–589 (2021).

37. Pettersen EF, et al. UCSF Chimera--a visualization system for exploratory research and. analysis. J Comput Chem 25, 1605–1612 (2004).

38. Emsley P, Cowtan K. Coot: model-building tools for molecular graphics. Acta Crystallogr D Biol Crystallogr 60, 2126–2132 (2004).

39. Afonine PV, et al. Real-space refinement in PHENIX for cryo-EM and crystallography. Acta Crystallogr D Struct Biol 74, 531–544 (2018).

40. Williams CJ, et al. MolProbity: More and better reference data for improved all-atom structure. validation. Protein Sci 27, 293–315 (2018).

41. Pettersen EF, et al. UCSF ChimeraX: Structure visualization for researchers, educators, and. developers. Protein Sci 30, 70–82 (2021).

