## Supplemental Data for "Structural basis of Retron-Eco8-mediated anti-phage defense"

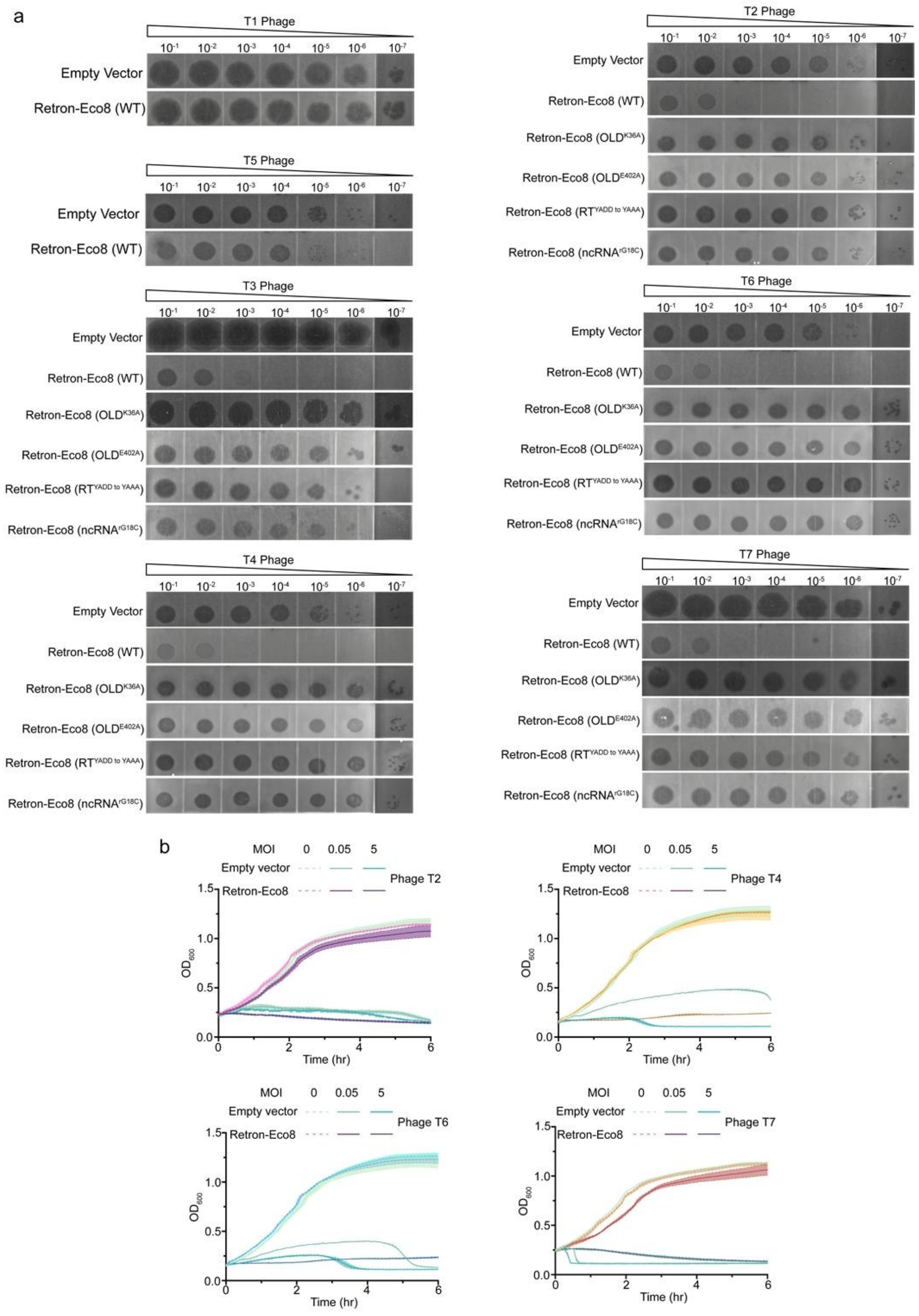


**Figure S1. The anti-phage activities of Retron-Eco8 system against various phages. Empty vector is used as the negative control.** (a) Plating assays of various phages on Retron-Eco8 system. (b) Phage infection in liquid cultures of the *E. coli* BL21(DE3) strain containing Retron-Eco8. Cells are infected at various MOI values. For each MOI, results of three experiments are presented as the average of three replicates with shaded areas indicating SD.

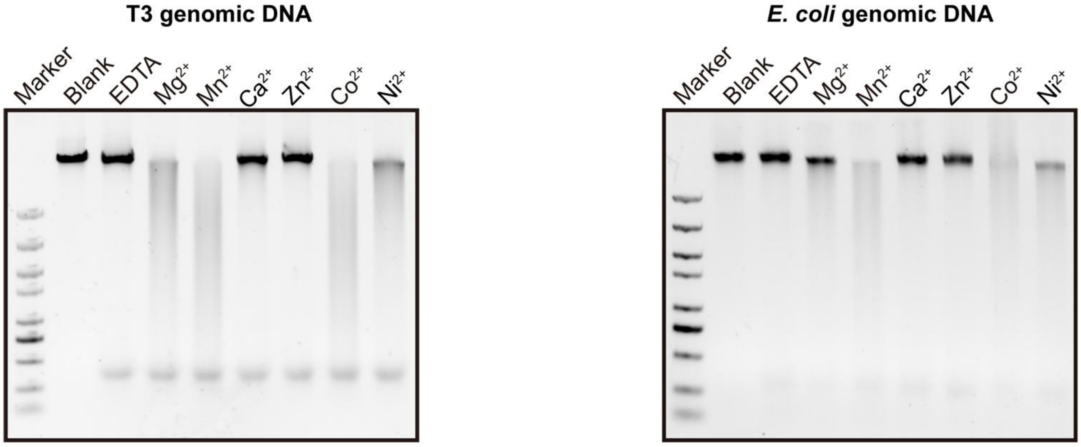


**Figure S2. The metal dependent nuclease activity of Retron-Eco8 against T3 phage and *E. coli* genomic DNA.**

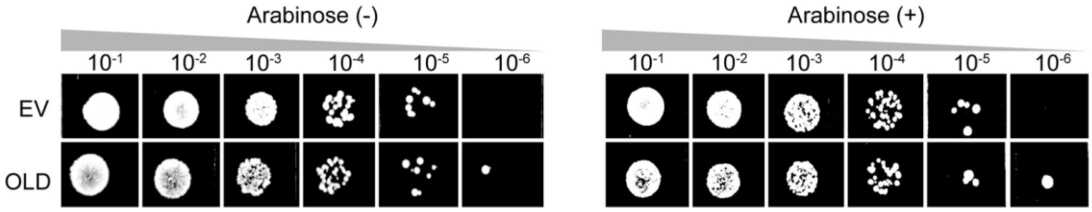


**Figure S3. Over-expression of OLD alone does not demonstrate cytotoxicity.**

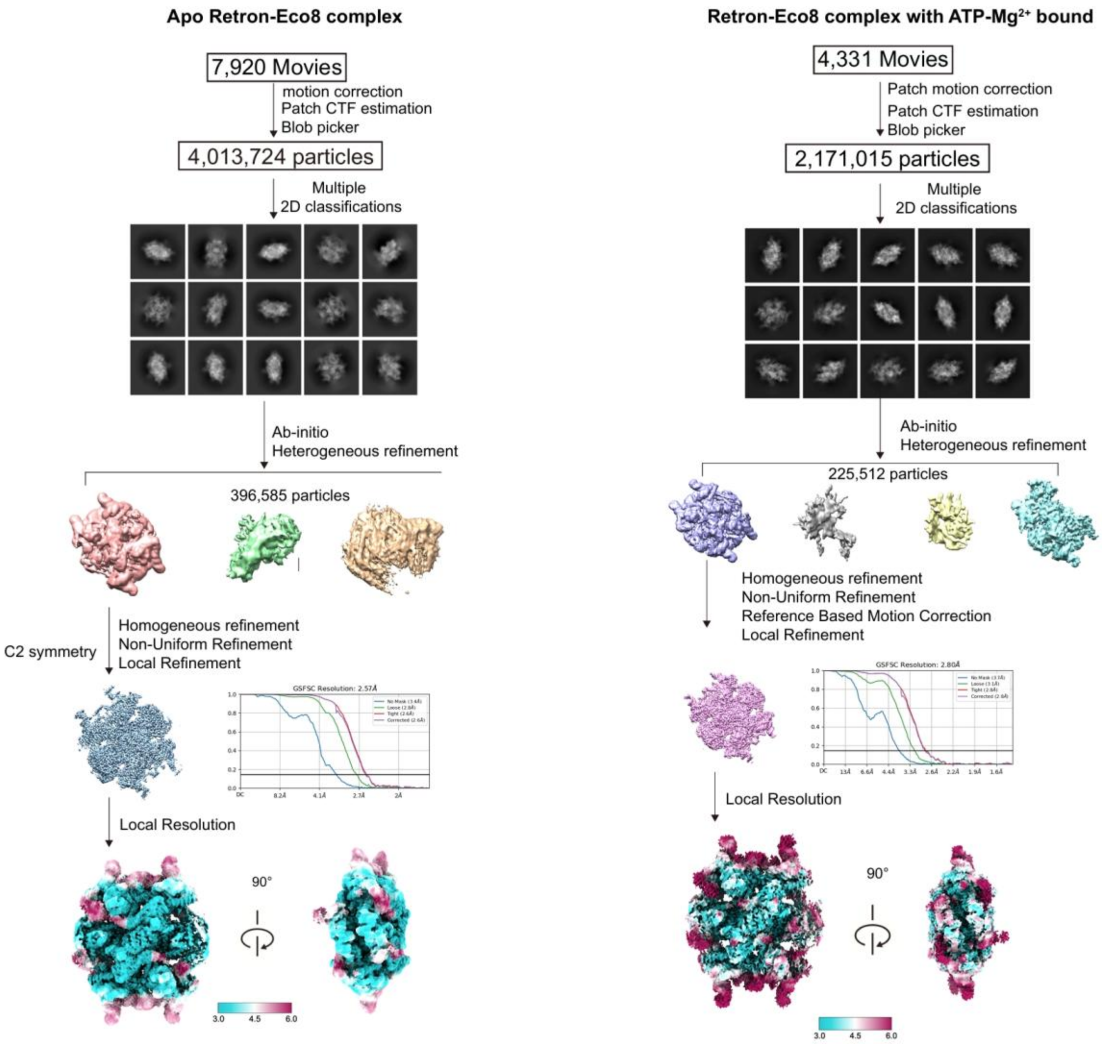


**Figure S4. Single particle cryo-EM analysis of Retron-Eco8 and Retron-Eco8 complexed with ATP–$Mg^{2+}$.** Representative cryo-EM micrographs, representative reference-free 2D-class averages, and data-processing workflows for Retron-Eco8 in the indicated states.

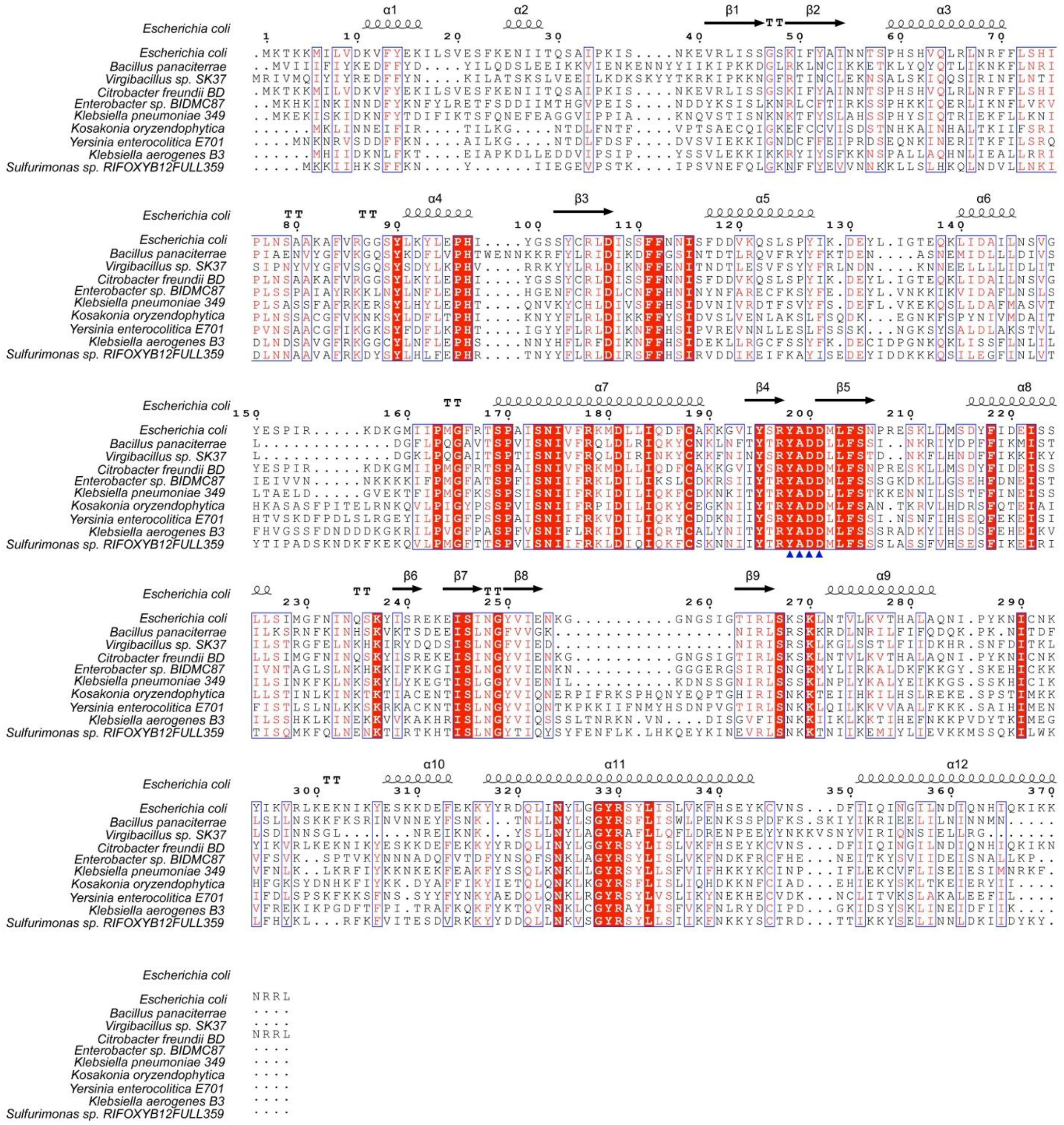


**Figure S5. Sequence alignment of Retron-Eco8 RTs of various species.** The secondary structure elements of *E. coli* Retron-Eco8 RT are shown above the alignment. The catalytic YADD motif is indicated by blue triangles.

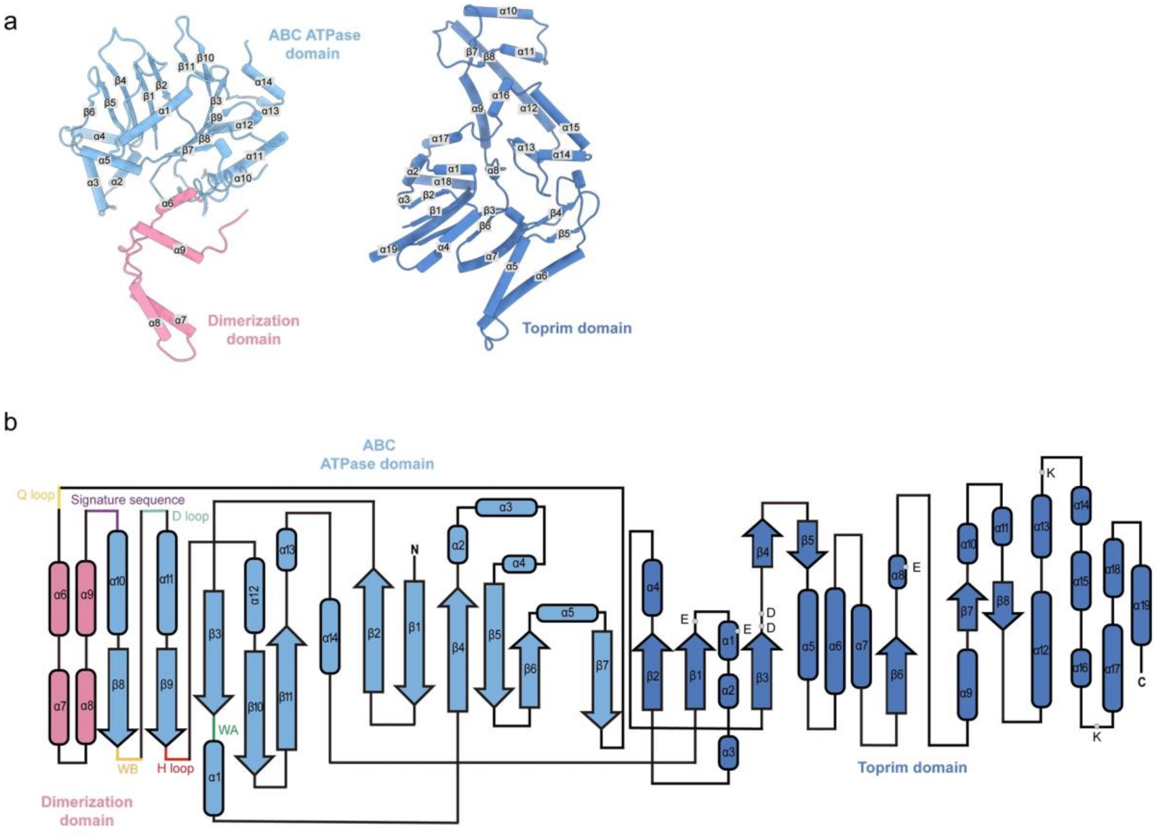


**Figure S6. Architecture of Retron-Eco8 OLD.** (a) The secondary structures of the ATPase, Dimerization and Toprim domains of Retron-Eco8 OLD. (b) Topology diagrams of Retron-Eco8 OLD. Six conserved motifs in the ATPase domain and the key residues in the Toprim nuclease active are labelled.

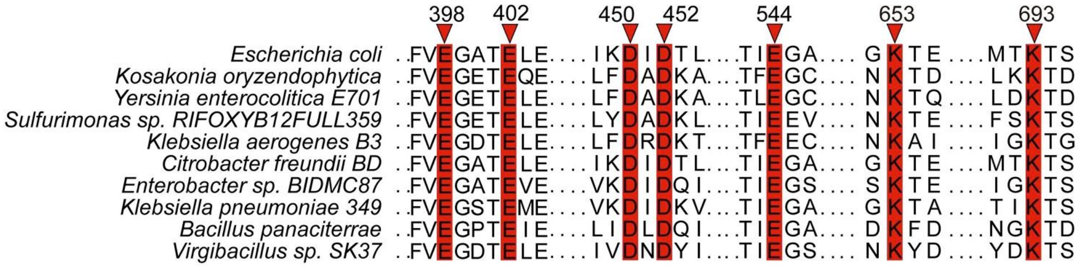


**Figure S7. Sequence alignment reveals that the key residues in Toprim domain active site are conserved among various species.**

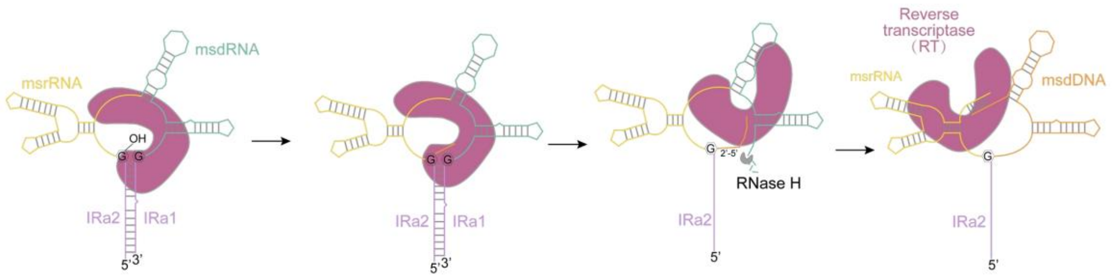


**Figure S8. Schematic depiction of the retron msDNA synthesis process using Retron-Eco8 as an example.**

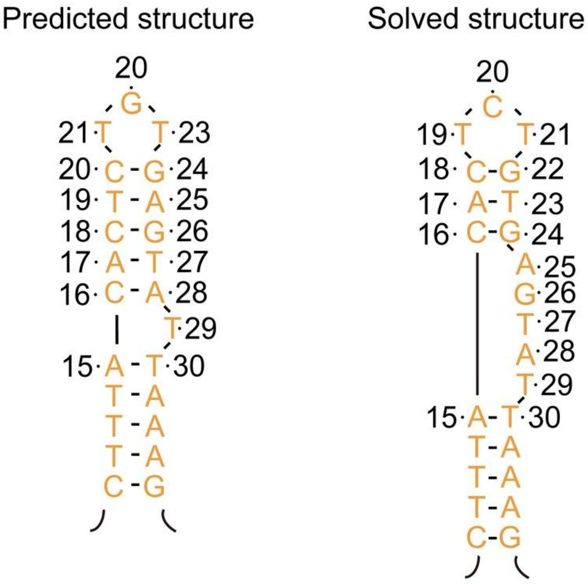


**Figure S9. Comparison of the predicted and solved DSLa structures of Retron-Eco8 msDNA.**

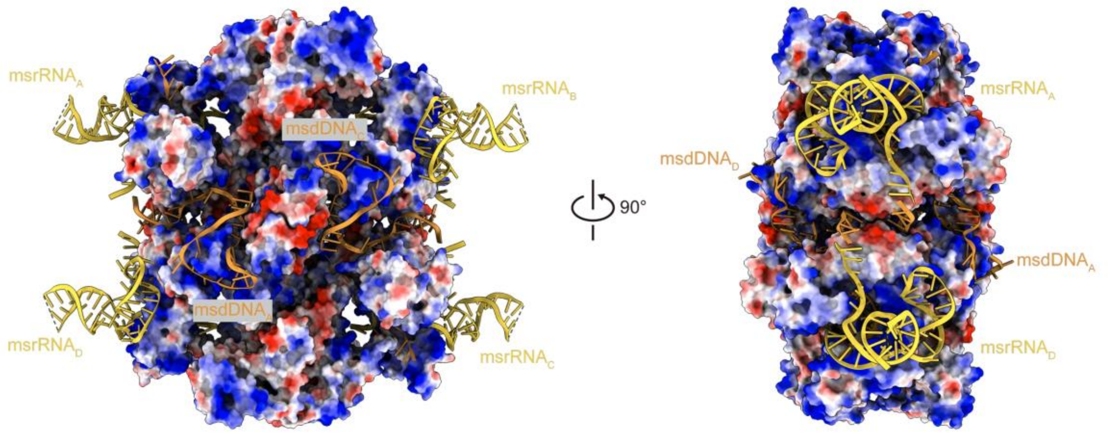


**Figure S10. Retron-Eco8 msDNA wraps the positively charged surfaces of RT and OLD. The electrostatic surface potential of Retron-Eco8 RT and OLD is shown. Blue and red (+/-5 kT/e) indicate the positively and negatively charged areas, respectively.**

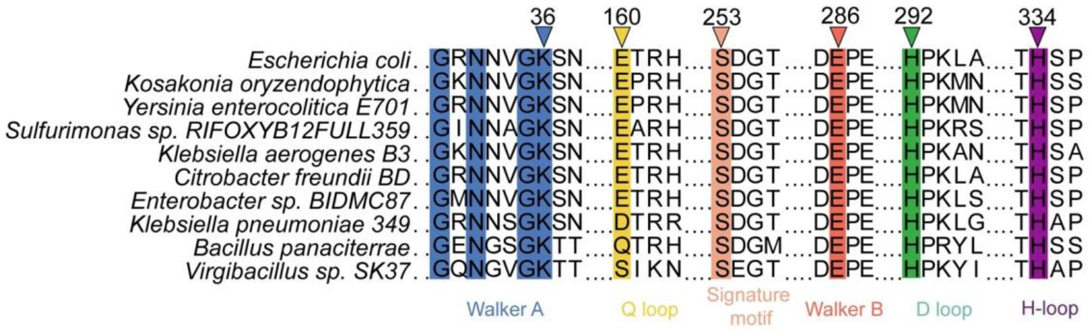


**Figure S11. Sequence alignment of Retron-Eco8 OLD ATPase domain of various species demonstrates the conserved key residues in ATPase active site.**

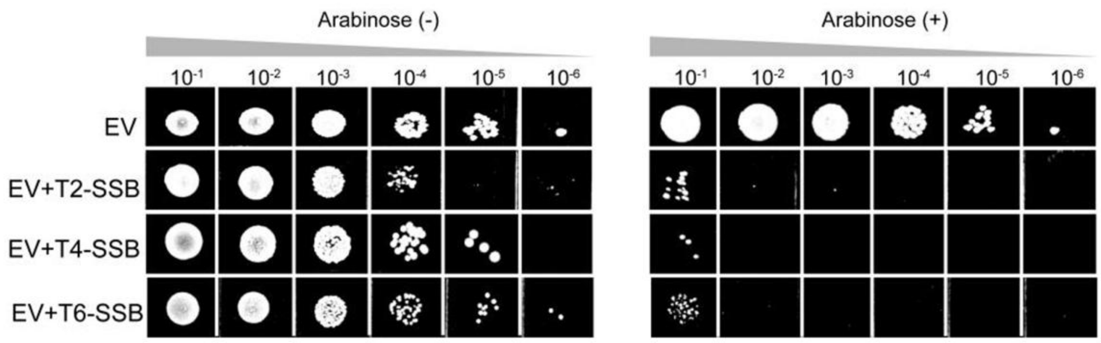


**Figure S12. Representative plating assay shows that over-expression of phage SSB proteins induces cytotoxicity. Empty vector (EV) is used as the negative control.**

**Table S1. Cryo-EM data collection, refinement, and validation statistics**

| Structure | Retron-Eco8 | Retron-Eco8 (ATP-$Mg^{2+}$) |
|---|---|---|
| EMDB ID | 66659 | 66663 |
| PDB ID | 9X94 | 9X9B |
| **Data collection and procession** | | |
| Magnification | 165000 | 165000 |
| Voltage (kV) | 300 | 300 |
| Electron exposure ($e^-/Å^2$) | 56.58 | 50 |
| Defocus range (μm) | -1.1~-1.8 | -0.8~-1.4 |
| Pixel size (Å) | 0.85 | 0.75 |
| Symmetry imposed | C2 | C1 |
| Initial particle images (no.) | 4,013,724 | 2,171,015 |
| Final particle images (no.) | 3,030,229 | 123,608 |
| Map resolution (Å) | 2.57 | 2.8 |
| FSC threshold | 0.143 | 0.143 |
| Map resolution range (Å) | 2.213-24.25 | 2.418-8.141 |
| Map sharpening *B* factor ($Å^2$) | -74.2 | -63.6 |
| **Refinement** | | |
| Initial model used (Protein) | Alphafold3 | Alphafold3 |
| Initial model used (Nucleotides) | de novo | de novo |
| Model resolution (Å) | 2.8 | 3.0 |
| FSC threshold | 0.5 | 0.5 |
| **Model composition** | | |
| Non-hydrogen atom | 44,906 | 45,453 |
| Protein residues | 4,396 | 4,445 |
| Nucleotide | 422 | 424 |
| Ligand | - | 188 |
| **B-factor ($Å^2$)** | | |
| Protein | 42.65 | 71.43 |
| Nucleotide | 60.37 | 90.50 |
| Ligand | | 56.39 |
| **R.m.s. deviations** | | |
| Bond lengths (Å) | 0.008 | 0.006 |
| Bond angles (°) | 0.827 | 0.736 |
| **Validation** | | |
| MolProbity score | 1.70 | 1.87 |
| Clashscore | 6.55 | 6.39 |
| Poor rotamers (%) | 0.46 | 0.12 |
| **Ramachandran plot** | | |
| Favored (%) | 95.15 | 90.81 |
| Allowed (%) | 4.83 | 9.19 |
| Outliers (%) | 0.02 | 0.00 |
